# DNAJB6 Isoform Specific Knockdown: Therapeutic Potential for LGMDD1

**DOI:** 10.1101/2022.11.17.516920

**Authors:** Andrew R. Findlay, May M. Paing, Jil A. Daw, Rocio Bengoechea, Sara K. Pittman, Shan Li, Feng Wang, Timothy M. Miller, Heather L. True, Tsui-Fen Chou, Conrad C. Weihl

## Abstract

Dominant missense mutations in DNAJB6, an HSP40 co-chaperone, cause limb girdle muscular dystrophy (LGMD) D1. No treatments are currently available. Two isoforms exist, DNAJB6a and DNAJB6b, each with distinct localizations in muscle. Mutations reside in both isoforms, yet evidence suggests only DNAJB6b is responsible for disease pathogenesis. Mechanistic data supports either a toxic gain of function, a dominant negative mechanism, or a combination of both. Knockdown treatment strategies involving both isoforms carry risk as DNAJB6 knockout is embryonic lethal. We therefore developed an isoform specific knockdown approach using morpholinos. Selective reduction of each isoform was achieved *in-vitro* in primary mouse myotubes and human myoblasts, as well as *in-vivo* in mouse skeletal muscle. To assess isoform specific knockdown in LGMDD1, we created primary myotube cultures from a *knock-in* LGMDD1 mouse model. Using mass spectrometry, we identified an LGMDD1 protein signature related to protein homeostasis and myofibril structure. Selective reduction of DNAJB6b levels in LGMDD1 myotubes corrected much of the proteomic disease signature towards wild type levels. While additional *in-vivo* functional data is required, these findings suggest selective reduction of DNAJB6b may be a viable therapeutic target for LGMDD1.

## Introduction

Limb girdle muscular dystrophy type-D1 (LGMDD1) is due to dominantly inherited mutations in *DNAJB6*. LGMDD1 patients have a wide range of age of onset and may have either a proximal or distal predominant pattern of weakness. Muscle biopsies demonstrate a vacuolar myopathy with aggregates as well as myofibrillar abnormalities involving the Z disc. No treatments are available. Standard of care is supportive in nature. Since its discovery in 2012, mutations in *DNAJB6* are the most common cause of dominantly inherited LGMD.^1^

DNAJB6 is a ubiquitously transcribed HSP40 co-chaperone with a wide array of functions. It acts as a tumor suppressor, prevents aggregation of polyglutamine containing proteins, and plays a role in viral replication cycles, just to name a few.^2^ Absence of *DNAJB6* is embryonic lethal due to aggregation of client proteins, keratin 8 / 18, within chorionic trophoblast cells, causing failure of chorioallantoic attachment during placental development.^3^ DNAJB6’s roles in muscle are thought to be related to protein homeostasis, but how dominantly inherited mutations in a ubiquitously expressed co-chaperone cause selective pathology in skeletal muscle is unknown.

DNAJB6 has two key isoforms, DNAJB6a a large nuclear predominant isoform, as well as DNAJB6b, a shorter isoform that localizes to sarcomeric structures in muscle, a key site of pathology seen in muscle biopsies from patients.^4^ Mutations associated with LGMDD1 are located in regions of *DNAJB6* that affect both isoforms. However, several lines of evidence suggest the B isoform preferentially contributes to disease pathogenesis.^4,5^ Zebrafish injected with mutant human DNAJB6b mRNA resulted in myofibrillar abnormalities, whereas mutant DNAJB6a mRNA did not.^5^ Additionally, transgenic mice overexpressing mutant human DNAJB6b in muscle caused a myopathy, whereas mice overexpressing mutant DNAJB6a were not different from controls.^4^ However, there is some evidence suggesting DNAJB6a is not completely dispensable in muscle, as a patient with a myofibrillar myopathy was found to have recessively inherited truncating frameshift mutations selectively affecting DNAJB6a.^6^ Overall, each isoform’s role in muscle and their contribution to LGMDD1 pathogenesis remains unclear. Additionally, there is mixed evidence whether disease pathogenesis occurs via a dominant negative effect, a toxic gain of function, or possibly a combination of both.4,5,7–11

Although gene knockdown is a common approach for treating dominantly inherited disorders, this approach may be deleterious in LGMDD1 based on DNAJB6 knockout data in cells and mice. However, haploinsufficiency may be tolerated, as heterozygous knockout mice have no reported phenotype, and human mutation databases such as gnomAD, contain frameshift and nonsense mutations scattered throughout DNAJB6 in presumably healthy patients.^3^ Recently, DNAJB6 haploinsufficiency was associated with a sick sinus syndrome phenotype in mice with no structural cardiac changes.^12^ Therapeutic approaches will therefore need to address the dominant mechanism of DNAJB6 mutations, but avoid potential deleterious effects of complete knockdown. Given the B isoform appears to preferentially contribute to disease pathogenesis, and the possibility of a dominant negative or toxic gain of function mechanism in LGMDD1, selective reduction of DNAJB6b levels could be a potential therapeutic target. In this work we designed morpholinos, a type of antisense oligonucleotide (ASO), to achieve DNAJB6 isoform specific knockdown and assess the therapeutic potential of selective DNAJB6b reduction in LGMDD1.

## Results

The mechanism governing DNAJB6 isoform expression was recently characterized and involves a competition between alternative polyadenylation and alternative splicing (Fig 1A).^13^ Production of the long A isoform depends on activation of intron 8 splicing and use of a strong distal polyadenylation signal (PAS), whereas production of the short B isoform depends on a lack of intron 8 splicing and use of a weak proximal PAS.

**Figure 1.**
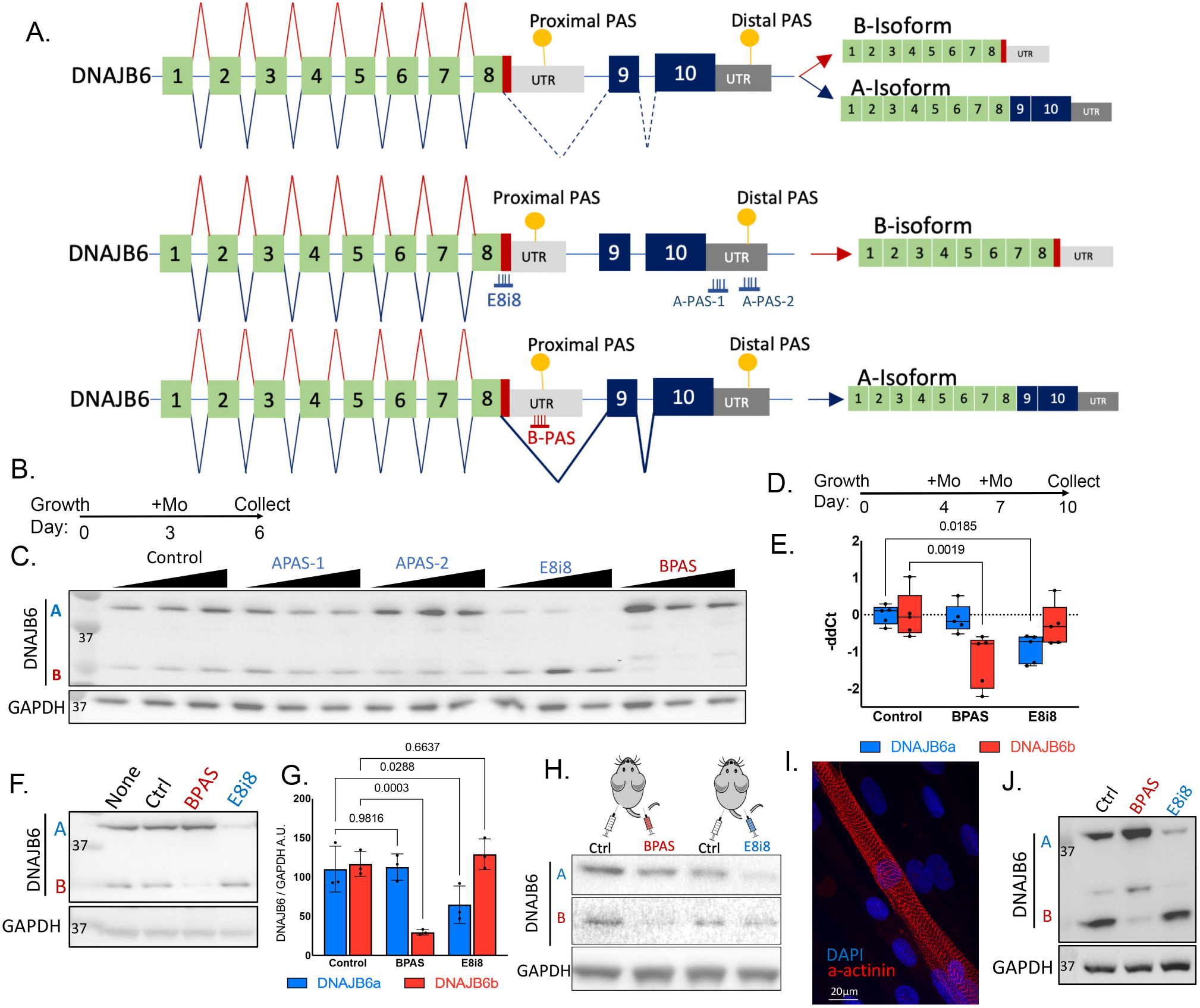
Selective DNAJB6 isoform knockdown *in-vitro* in primary mouse myotubes and primary human myoblasts and *in-vivo* in mouse skeletal muscle. (A) Map of DNAJB6 gene structure and transcripts generated by competition between alternative splicing and polyadenylation. Morpholinos designed to selectively reduce DNAJB6b transcript (BPAS) targets the proximal PAS, sterically blocking this sequence, in theory, preventing its polyadenylation. Morpholinos designed to selectively reduce DNAJB6a transcript by targeting the distal PAS (A-PAS1, A-PAS2), or by blocking intron 8 splicing (E8i8). (B) Culture and treatment timeline for screening morpholinos (Mo) in primary mouse myotubes. (C) Western blot of lysates from primary mouse myotubes treated with morpholinos to achieve isoform specific knockdown. BPAS and E8i8 appear to result in a dose dependent reduction of their intended targets. Morpholino doses: 2.5μM, 5μM, and 10μM. (D) Culture and treatment timeline for optimizing DNAJB6 isoform knockdown in mature myotubes. Morpholino dose: 5μM. (E) Boxplots of DNAJB6 isoform specific real-time quantitative PCR (RT-qPCR) of primary myotubes treated with control, BPAS, or E8i8 morpholinos. BPAS results in significant reduction of only the B isoform transcript, whereas E8i8 causes a significant reduction of only the A isoform transcript. Dots represent experimental replicates (n=5). Each experimental replicate was generated from 3 technical replicates. Whiskers represent the range, the upper and lower borders of the box represent the first (25%) and third quartiles (75%), and the horizontal line is the median. One-way ANOVA with Tukey’s post-hoc test comparing BPAS or E8i8 treatment to control. Only significant comparisons are shown. (F) Representative western blot of lysates from primary myotubes treated with optimized culture and morpholino treatment timeline. (G) Quantitation of western blot. Dots represent experimental replicates (n=3). Results are presented as mean +/-SD. One-way ANOVA with Tukey’s post-hoc test was used to compare BPAS or E8i8 treatment’s impact on DNAJB6a and DNAJB6b levels vs. control. (H) Western blot of skeletal muscle lysates from C57/B6 mice 4 days after intramuscular injection of BPAS or E8i8 into TA of 1 leg, and control morpholino in the contralateral leg. 10μg morpholino per leg. DNAJB6 membrane was split between DNAJB6a and DNAJB6b to develop separately due to the large excess of DNAJB6a relative to DNAJB6b in mature skeletal muscle. (I) Primary myoblasts isolated from LGMDD1 muscle biopsy (p.F89I), differentiated into myotubes, and stained for alpha-actinin (red) and nuclei (blue). Mature myotubes with well-formed sarcomeres are present. (J) Western blot of lysates from LGMDD1 primary human myoblasts treated with morpholinos for 3 days demonstrating effective isoform specific reduction with BPAS and E8i8.

We designed two strategies using morpholinos to prevent production of DNAJB6a: blocking DNAJB6a’s PAS, theoretically destabilizing DNAJB6a mRNA (A-PAS-1, A-PAS-2), or blocking intron 8 splicing (E8i8), to prevent DNAJB6a mRNA production.

DNAJB6b has no unique sequence compared to DNAJB6a at the DNA or pre-mRNA level. The only sequence unique to the DNAJB6b isoform occurs at the mRNA level, and includes 35 coding bases and the 3’ untranslated region (UTR) (Fig 1A). We therefore designed a morpholino (BPAS) targeting DNAJB6b’s PAS, theoretically impairing its polyadenylation and destabilizing the transcript, resulting in production of only the A isoform (Fig 1A)

Morpholinos designed for selective reduction of each DNAJB6 isoform were first screened using C57/B6 primary myoblasts differentiated into myotubes. Myoblasts were grown for 3 days to near confluency before media was changed and morpholinos were added (Fig 1B). Morpholinos were taken up by gymnosis due to the presence of an octaguanidine dendrimer moiety that improves uptake. Myotubes formed over the next 3 days and cells were collected for western blot (Fig 1C). The E8i8 morpholino effectively reduced the DNAJB6a isoform and BPAS selectively reduced the DNAJB6b isoform relative to myotubes treated with a control morpholino. Morpholinos targeting the A isoform’s PAS (A-PAS-1, A-PAS-2) were not effective (Fig 1C).

We optimized our protocol to maximize both isoform specific knockdown and myotube maturity (Fig 1D). Selective reduction of each isoform was detected at both the RNA and protein levels (Fig 1E-G). In certain instances, following E8i8 treatment we noted an increase in the B isoform, and following BPAS treatment we noted increased A isoform levels (Fig 1C).

We conducted preliminary *in-vivo* testing in C57/B6 mice using intramuscular (IM) tibialis anterior (TA) injections and collected muscle 4 days later. Mice received control morpholino in 1 leg and either BPAS or E8i8 in the other (Fig 1H). Western blot of skeletal muscle lysates demonstrated BPAS and E8i8 are capable of selective isoform reduction at the protein level *in-vivo* (Fig 1H).

To test feasibility in a human model of disease, we isolated primary myoblasts from skeletal muscle biopsies of LGMDD1 patients. These myoblasts readily differentiate into myotubes with sarcomeric structures (Fig 1I). Treatment of LGMDD1 myoblasts containing a p.F89I mutation with BPAS or E8i8 for 3 days also achieved the intended isoform specific reductions at the protein level (Fig 1J).

We next wondered how isoform specific knockdown might impact an LGMDD1 disease model. We previously created a *knock-in* LGMDD1 mouse model with a heterozygous p.F90I mutation (F90I+/-) (orthologous to human p.F89I).^7^ These mice develop a myopathic phenotype by at least 1 year of age.^7^ Primary myoblasts isolated and cultured from these mice had no difference in proliferation rate compared to WT primary mouse myoblasts (Fig 2A). We also detected no differences between WT and F90I+/-in their ability to form myotubes (fusion index) or their myofibrillar structure as shown by alpha actinin staining (Fig 2B,C). Although there were no obvious phenotypic abnormalities in the LGMDD1 primary mouse myotube cultures, this could be related to LGMDD1 being a degenerative disorder of mature skeletal muscle, not a condition with congenital onset related to dysfunctional muscle formation.

**Figure 2.**
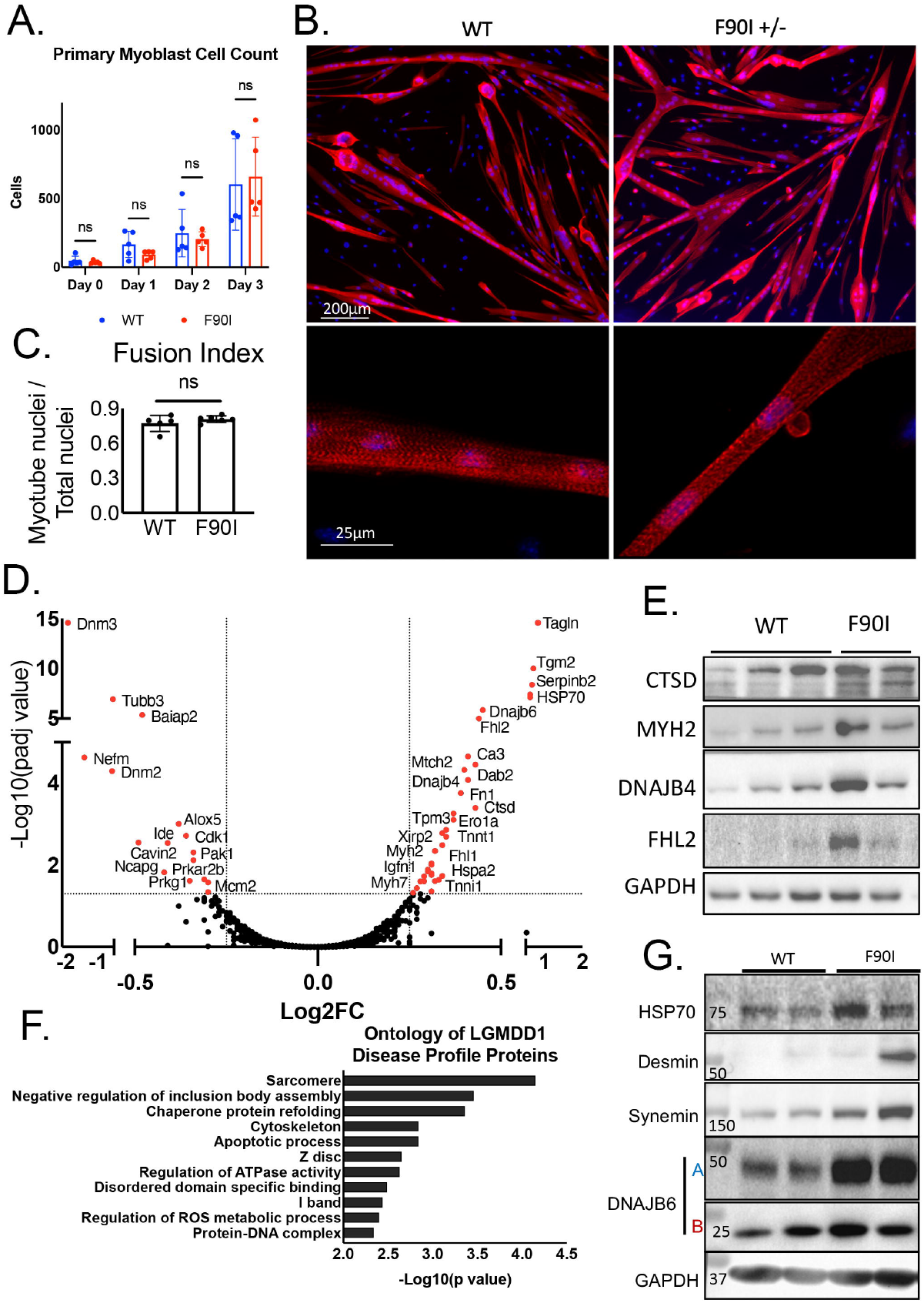
LGMDD1 proteomic signature identifies proteins involved in myofibrillar structure and protein homeostasis in primary mouse myotubes and skeletal muscle. (A) Quantitation of primary myoblast proliferation from F90I+/-and WT mice shows no significant differences. Myoblasts were plated at equal densities, collected at the indicated time points, and stained with DAPI. Results presented as mean +/-SD. Dots represent experimental replicates (n=5). Statistical comparisons made via 2-way ANOVA with Bonferroni post-hoc test. (B) Primary myotubes at day 3 of differentiation from F90I+/-and WT mice develop multinucleated myotubes. Bottom panels are higher magnification demonstrating well formed sarcomeric structures. Alpha-actinin (red) and DAPI (blue). (C)Quantitation of fusion index (myotube nuclei / total nuclei) shows no difference between WT and LGMDD1 cell lines. Results presented as mean +/-SD. Dots represent experimental replicates (n=5). 2-tailed, unpaired Student’s t-test. (D) Volcano plot of mass spectrometry data comparing F90I+/-to WT myotubes (n=3 experimental replicates). Log2FC ratio is F90I / WT. Proteins with significantly altered abundances are highlighted in red. (E) Western blot validation of several proteins identified in (D) with significantly altered abundances. Each lane is lysate from an experimental replicate of the mass spectrometry experiment. The third F90I+/-replicate is not shown due to absence of GAPDH signal. (F) Ontological analysis of proteins identified to have altered abundances in F90I+/-myotubes. (G) Western blot of skeletal muscle lysates (TA) from F90I +/-mice (n=2) and WT controls (n=2) at 2 months of age (same age at which primary myoblasts were isolated). Protein homeostasis proteins: DNAJB6 and HSP70. Myofibrillar proteins: desmin and synemin.

To better characterize a monitorable disease phenotype in LGMDD1 primary myotubes, we sought to use a broad “omic”-based approach as DNAJB6 has many functions, and it’s unclear which of those functions are impacted by disease mutations.^2^ We therefore decided to use mass spectrometry as chaperone dysfunction might be most directly captured at the protein level. By comparing with WT myotubes, we aimed to characterize a disease signature of altered protein abundances in the F90I+/-myotubes.

WT and F90I+/-myotube cultures were treated with control morpholinos starting at day 4 in culture, replenished at day 7, and collected at day 10 for tandem mass tag (TMT) labeling and mass spectrometry. Using stringent criteria (FDR<0.01, coverage >10%, peptides > 10) to ensure high confidence hits, we found 54 proteins with significantly altered abundances (Log2 fold change (Log2FC) >0.25 or <-0.25, padj <0.05) between WT and F90I +/-myotubes (Fig 2D,E, Table S1). We validated several of these changes via western blot (Fig 2E, 3C). Ontological analysis of these proteins supports their relevance to disease pathogenesis involving altered protein homeostasis and myofibrillar proteins (Fig 2F). Consistent with the *in-vitro* proteomic profile, we similarly found altered levels of proteins involved in myofibrillar structure and protein homeostasis in mature skeletal muscle from F90I+/-mice (Fig 2G).

DNAJB6b is known to localize to myofibrillar structures like the Z disc, whereas DNAJB6a localizes to myonuclei.^4^ Given the proteomic signature appeared more relevant to myofibrillar proteins than nuclear proteins, and because DNAJB6b is thought to preferentially impact LGMDD1 pathogenesis, we focused on BPAS’s potential impact on the proteomic signature identified in F90I+/-myotubes. Myotubes treated with BPAS morphologically appeared no different compared to those treated with control morpholinos (Fig 3A). Following F90I+/-myotube treatment for 6 days with BPAS, 31 of the 54 (57%) disease profile proteins returned back to WT levels (Fig 3B). Western blot validation of HSP70, a protein directly linked to LGMDD1 pathogenesis, as well as DNAJB6b, confirmed a return to WT levels (Fig 3C-E).^7^ Ontological analysis of proteins whose levels were corrected, again involved categories related to protein homeostasis and myofibrillar proteins, whereas ontological analysis of those not corrected by BPAS treatment included many categories with no currently known connection to LGMDD1 pathogenesis (Fig 3E,F). Given the therapeutic potential of selective B isoform reduction we evaluated for un-intended binding sites of BPAS using nucleotide BLAST. Within the mouse transcriptome, the closest off-target sites had at least 7 mismatches: *MAP3K3* (72%), whereas off-target sites in the human transcriptome had at least 9 mismatches: *ADGRG4* (64%).

**Figure 3.**
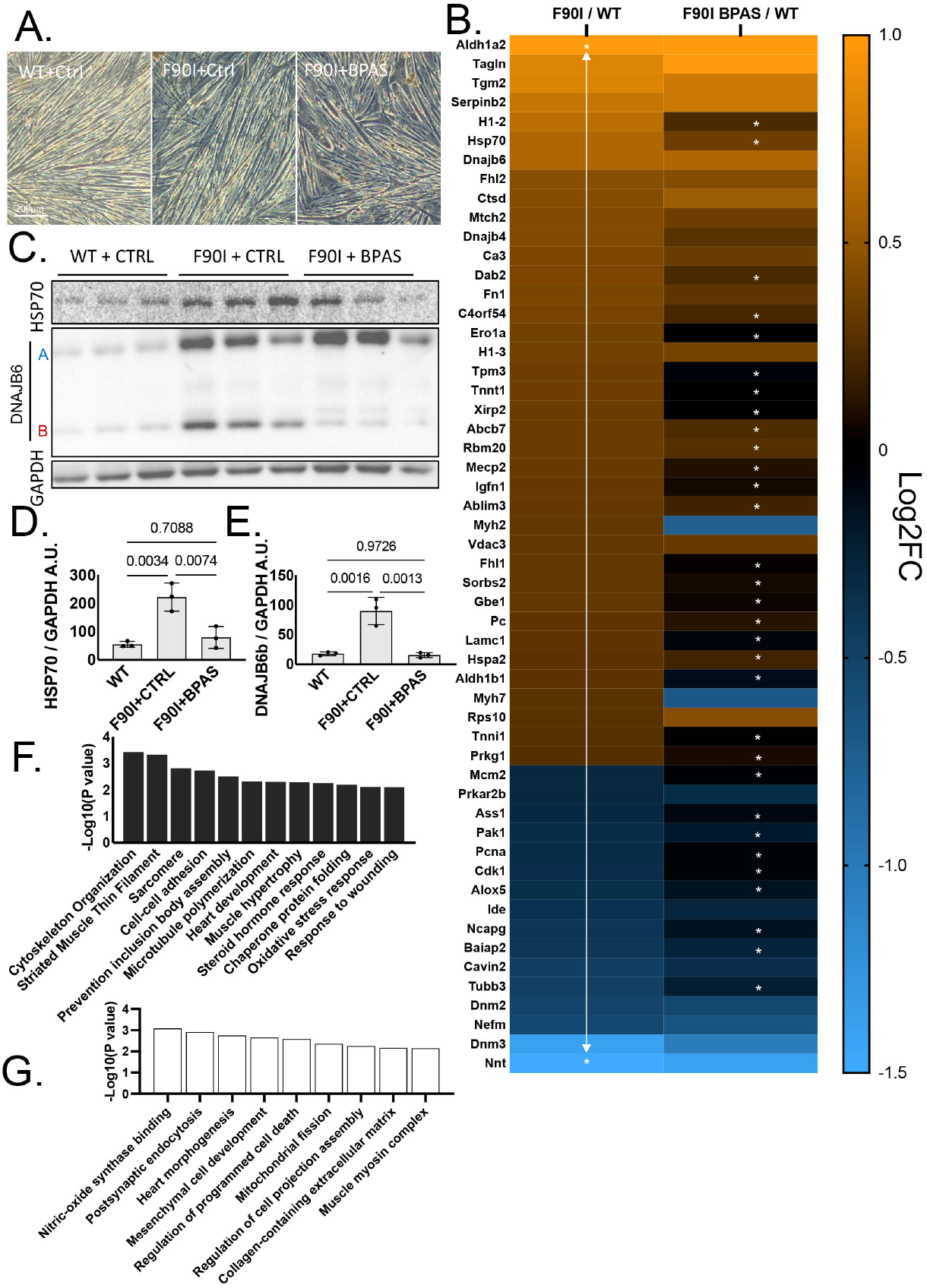
DNAJB6b isoform reduction normalizes proteomic signature in LGMDD1 primary mouse myotubes. (A) Brightfield images of myotubes treated with control and BPAS morpholinos demonstrate no morphological changes. (B) Heatmap of 54 proteins with altered abundance in F90+/-myotubes. Data expressed as log 2-fold change relative to WT control treated myotubes. Higher abundance is indicated by orange, less abundance is represented by blue, and no change is black. Following BPAS treatment of F90I+/-myotubes for 6 days, 31/54 (57%) proteins returned to WT levels. Asterisks denote the 31 proteins whose relative abundances were restored to WT levels (no significant difference compared to WT levels). Data generated from 3 experimental replicates. (C) Western blot of myotube lysates from mass spectrometry experiment, validating HSP70 normalization following DNAJB6b reduction. (D) Quantitation of HSP70 western blot. Data presented as mean +/-SD. Dots represent experimental replicates (n=3) from 3C. (E) Quantitation of DNAJB6b western blot. Data presented as mean +/-SD. Dots represent experimental replicates (n=3) from 3C. (F) Ontological analysis of 31 proteins corrected to WT levels with BPAS treatment. (G) Ontological analysis of proteins not corrected to WT levels with BPAS treatment.

## Discussion

Since 2012 when mutations in *DNAJB6* were identified as a cause for dominantly inherited LGMD, there has been tremendous progress in the LGMDD1 field. Numerous cell and animal models have been generated, promising therapeutic targets are being identified, and outcome measures for future clinical trials are being defined.^4,7,8,10^ Despite this progress, there are no treatments for LGMDD1, and therapeutic target identification is still a significant need.

Our previous studies and others have suggested the short B isoform of DNAJB6 preferentially contributes to LGMDD1 pathogenesis.^4,5^ This is supported by its localization to myofibrillar structures within skeletal muscle, a key site of pathology seen in human biopsies.^5,14,15^ Although the disease mechanism may involve a dominant negative, toxic, or gain of function mechanism, absence of both DNAJB6 isoforms is known to be embryonic lethal in mice.^3^ While it’s unknown if postnatal absence of DNAJB6 is deleterious *in-vivo*, therapeutic knockdown approaches for LGMDD1 that avoid complete DNAJB6 knockdown may have a wider therapeutic index.

Here we show the feasibility of DNAJB6 isoform specific reduction and the potential of using morpholinos for LGMDD1 therapy via DNAJB6b knockdown. Our choice to use morpholinos was influenced by the significant amount of preclinical and clinical studies using morpholinos in neuromuscular disorders. Antisense oligonucleotide (ASO)-based splice-modifying therapies for Duchenne muscular dystrophy (DMD) and spinal muscular atrophy have created a model framework for moving this class of drugs towards the clinic. Our study used morpholinos modified with an octaguanidine dendrimer to improve cellular uptake *in-vitro* and *in-vivo*. Although ideal for pre-clinical proof of principle studies, this form of morpholino has not been studied in human trials. Additionally, there are safety concerns related to fatal clotting episodes with this form of morpholino in mice.^16^ Future pre-clinical studies will use a more clinically translatable chemistry such as next-generation peptide-conjugated phosphorodiamidate morpholino oligomers (PMO), PMOs conjugated to TfR1-targeting antigen-binding fragments (Fab), or antibody-oligonucleotide conjugates (AOCs). Another more translatable approach to achieve DNAJB6b isoform knockdown, with the added benefit of a single administration potentially providing lifelong treatment, is an adeno-associated virus (AAV) approach to deliver U7-snRNA targeting the proximal PAS of DNAJB6b. AAV delivered U7-snRNAs are currently in clinical trials for DMD as an exon skipping therapy for exon 2 duplications, and have been used pre-clinically in fascioscapulohumeral muscular dystrophy (FSHD) to reduce DUX4 expression by similarly targeting the PAS.^17,18^

Aside from showing reduction of DNAJB6 isoform levels at the RNA and protein levels in primary mouse myotube cultures and primary LGMDD1 patient derived myoblasts, we demonstrated selective knockdown of DNAJB6 isoform protein levels in the TA muscle of mice with IM injections. We also provided preliminary evidence of safety against potential off-target effects. Nucleotide BLAST results using BPAS’s sequence against the mouse and human transcriptomes yielded no significant hits with the closest targets having at least 10 mismatches (60% sequence complementarity).

Previous studies characterizing the mechanism regulating DNAJB6 isoform expression similarly used morpholinos to modify DNAJB6 isoform expression, but focused on reducing the A isoform, using a morpholino similar to E8i8.^13^ DNAJB6a is known to play a role in the replication cycle of several viruses, including human immunodeficiency virus (HIV).^19^ It was therefore hypothesized reduction of DNAJB6a could be a target for a broad-spectrum antiviral agent. Selective DNAJB6a reduction did in fact impair HIV and respiratory syncytial virus (RSV) replication in human cell lines without any apparent negative effects.^13^ However, a patient with a myofibrillar myopathy was recently reported to have homozygous recessive frameshift mutations selectively affecting the DNAJB6a isoform, resulting in loss of A isoform protein expression.^6^ These findings raise concern about the safety of any intervention that greatly reduces DNAJB6a levels. We occasionally noticed an increase in DNAJB6a levels following treatment with BPAS (Fig 1C, 1J, 3C). While the mechanism for this observation remains unclear, it is possible that by blocking the B isoform’s PAS, the stronger distal PAS is preferentially utilized, shunting DNAJB6 mRNA production towards the A isoform. Future research will need to evaluate any risks associated with increased DNAJB6a levels, especially given its association with viral replication cycles and its role as a tumor suppressor.^13,20–23^

Although the BPAS morpholino significantly reduced DNAJB6b levels, other sequences targeting DNAJB6’s proximal PAS may be more effective. Future work will aim to optimize this antisense sequence targeting this region. While our culture system of primary *knock-in* LGMDD1 myotubes does not display any overt morphological abnormalities, they do have a distinct, correctable, proteomic signature which will be useful for screening future therapeutics *in-vitro*. Similar omics-based approaches are now commonly used to capture an unbiased disease signature, even in mild or pre-symptomatic disease stages, and then leveraged as a therapeutic biomarker.^24–27^ Our results are encouraging as they suggest DNAJB6b reduction is beneficial in reversing the molecular effects of mutant DNAJB6 and improving a pathological *in-vitro* phenotype.

In conclusion, using primary LGMDD1 mouse myotubes, primary human LGMDD1 myoblasts, and skeletal muscle from a *knock-in* LGMDD1 mouse model, we were able to show that our morpholinos could selectively reduce DNAJB6 isoforms. The functional effects of DNAJB6 isoform reduction on mature skeletal muscle remains to be determined with a systemic treatment study. These studies are ongoing and will complement our *in-vitro* findings which show a limited but promising view of the potential for DNAJB6b isoform reduction as a therapeutic target for LGMDD1. Additionally, we outline the use of an *in-vitro* therapeutically correctable proteomic disease signature in LGMDD1 myotubes. This pharmaco-responsive signature of altered protein homeostasis and myofibrillar proteins extrapolates *in-vivo* to our LGMDD1 mouse model. Taken together, we expect these preliminary signs of therapeutic benefit from DNAJB6b isoform reduction, and the unbiased proteomic method for screening therapeutics in this study to facilitate progress in the field toward production of viable treatments for LGMDD1.

## Materials & methods

### Morpholino Treatment

Octaguanidine dendrimer-conjugated morpholino oligonucleotides (Gene Tools) were used in this study. BPAS (5’-TGCACCAAACACATTCGCATTTATT-3’; Gene Tools), E8i8 (5’-CGTAGCAGGTGCTCCTTACCATTTA-3’; Gene Tools), A-PAS-1 (5’-ACATTTTATTTACGTCAGTAGCACT -3’), A-PAS-2 (5’-AGACCTGCTTTTATTTTTCATAGTA-3’), and negative control (5’-CCTCTTACCTCAGTTACAATTTATA-3’; Gene Tools). Morpholinos were added to cell culture media for the indicated time periods for each experiment. *In-vitro* screening of morpholinos in primary mouse myotubes used doses of 2.5μM, 5μM, and 10μM. BPAS, E8i8 and control morpholinos were subsequently used at 5μM for treatment of primary mouse myotubes and primary human myoblasts. For *in-vivo* treatments, after oxygen exposure, mice were anesthetized with isoflurane and injected intramuscularly (TA) with morpholinos (10μg morpholino diluted in 20μl phosphate buffered saline [PBS, Gibco]).

### Primary mouse myoblast isolation and culture

Primary mouse myoblasts were isolated and cultured as previously described.^28^ At 2 months of age, WT and F90I+/-mice were sacrificed, hind limb muscles were isolated, and excess connective tissues and fat were cleaned in sterile PBS. Muscle tissues were minced into small pieces with scissors and added to a 50mL conical tube with 1mL of enzyme mix (500 U/mL type II collagenase [MP biomed], 1.5U/mL Collagenase D [Sigma] 2.5U/mL Dispase II [Sigma], 2.5mM CaCl_2_, PBS up to 1mL). The mixture was shook at 37°C for 1 hour, pelleted at 2000rpm for 5 minutes, and the supernatant was aspirated. The pellet was resuspend in proliferation medium (high-glucose Dulbecco’s Modified Eagle Medium [DMEM, Gibco], supplemented with 20% fetal bovine serum, [FBS, Gibco], 10% horse serum [HS, Remel], 0.5% chicken embryo extract [CEE, Accurate Chemical and Scientific], 2.5 ng/mL basic fibroblast growth factor [bFGF, PeproTech], 10μg/mL Gentamycin, 1% Antibiotic/Antimycotic [AA, Gibco], and 2.5μg/mL Plasmocin prophylactic [Invivogen, San Diego, CA]), seeded on 10% Matrigel-coated dishes (45 μg/cm^2^) at 10-20% surface coverage, and incubated at 37°C and 10% CO_2_. After cellular outgrowth from muscle explants was observed, tissues were transferred to new Matrigel coated plates for additional rounds of myoblast outgrowth. Prior to confluence of cellular outgrowth, cells were detached using 0.25% trypsin and to improve myoblast purity, cells were pre-plated on collagen coated plates (type I rat tail collagen [CORNING] in sterile water at 5 μg/cm^2^) in growth medium, for 1 hour at 37°C and 10% CO_2_. The supernatant was then transferred to Matrigel-coated dishes. Myoblasts were subsequently grown on Matrigel-coated dishes in proliferation medium, while cells on the collagen coated plates were discarded. For myoblast differentiation, switching to a low serum media resulted in rapid myotube formation, contractions, and subsequent detachment. To avoid rapid detachment and facilitate slower differentiation and longer treatment with morpholinos, we plated myoblasts at a high baseline confluency in growth media, and infrequently changed the media to allow serum levels to be gradually consumed as myoblasts proliferate or differentiate into myotubes. Timing of media changes is detailed for each experiment.

### Primary human myoblast isolation and culture

Healthy individuals and individuals with genetically confirmed LGMDD1 were biopsied as part of ongoing research studies as approved by the Institutional Review Board of Washington University St. Louis (IRB# 202102146). Written informed consent was obtained according to the Declaration of Helsinki from all participants or a parent/legal guardian. Biopsies were performed at Washington University St. Louis by co-authors ARF and CCW. TA core biopsies were obtained using sterile technique and equipment throughout the procedure. The skin overlying the biopsy site was marked and anesthetized with lidocaine. The co-axial introducer needle was inserted in the middle of the muscle belly. Three samples, totaling 45 grams were collected from the TA with a micro-biopsy needle (Argon SuperCore™ Semi-Automatic Biopsy Instrument 14 GÅ∼9 cm, with 13 GÅ∼3.9 cm co-axial needle). After each pass, the semi-automatic biopsy instrument was removed from the co-axial introducer needle. Muscle samples were carefully removed and placed into vials with collection medium (Hanks’ Balanced Salt Solution [HBSS, Gibco], 1% Penicillin-Streptomycin (PS), 0.25ug/mL amphotericin B).

Myoblasts were then isolated and cultured as previously described with minor modifications.^28,29^ Muscle was minced into small pieces using a McIlwain tissue chopper (Mickle Laboratory Engineering, Surrey, UK), added to a 15mL conical tube with 1mL of enzyme mix (1.5U/mL Collagenase D, 2.5U/mL Dispase II, 2.5mM CaCl_2_, HBSS up to 1mL), and shook at 37°C for 1 hour. Tissue fragments were pelleted at 2000 rpm for 5 minutes and the supernatant was aspirated. The pellet was resuspended in human myoblast proliferation medium (20%FBS, 10% HS, 0.5% CEE, 2.5ng/mL bFGF, 1% penicillin-streptomycin, 0.25 ug/mL amphotericin), plated on 10% Matrigel coated plates, and incubated at 37°C and 10% CO_2_. After 2-3 days, proliferation medium was replaced. Approximately 5 days later, when the muscle tissue is adherent and myoblast outgrowth is observed, but before cells become confluent, cells were detached using TrypLE express [Gibco], and pre-plated on collagen-coated plates for 1 hour at 37°C at 10% CO_2_. The supernatant was then transferred to 10% Matrigel-coated dishes (45 μg/cm^2^) in human myoblast proliferation medium. Additional rounds of pre-plating (typically 2 rounds total) were performed as needed to enrich for cells with myoblast morphology. Myoblasts are grown in human myoblast proliferation medium until 85-95% confluency before switching to differentiation media (DMEM, 5% HS, 1% PS, 0.25μg/mL amphotericin B) to form myotubes. Medium is changed every 24-36 hours.

### Animal and Experimental Protocols

All animal experimental protocols were approved by the Animal Studies Committee of Washington University School of Medicine. CRISPR/Cas9-mediated *knock-in* F90I +/-mice were generated as previously described, by the mouse genetics core facility for transgenic animal production at Washington University.^7^ Animal lines were bred on a C57/B6 background (The Jackson Laboratory) to at least the F5 generation. Control animals were F90I -/-littermates. Mice were housed in a temperature-controlled environment with 12-hour light/12-hour dark cycles, in which they received food and water ad libitum. Mice were euthanized, and skeletal muscle was dissected. For western blot, muscle was flash frozen in liquid nitrogen and stored at −80°C.

### Immunofluorescence

Primary mouse myoblasts and myotubes were grown, stained, and imaged directly on cell culture plastic. Primary human myotubes were grown, stained, and imaged on glass coverslips coated with 0.1% gelatin. Cells were washed 3 times with PBS, fixed in 4% PFA for 10 minutes, permeabilized with 0.1% Triton X-100 in PBS for 10 minutes, and then blocked with 3% BSA in PBS for 30 minutes to 1 hour at room temperature. Cells were stained with primary antibody (anti-rabbit alpha-actinin) at 4°C overnight, followed by washing 3 times with PBS. Cells were incubated with Alexa Fluor 555-conjugated secondary antibody at room temperature for 1 hour and mounted with Mowiol media containing 4′,6-diamidino-2-phenylindole (DAPI).

Fusion index was determined as a ratio of nuclei number within multinucleated alpha-actinin positive myotubes to the total number of nuclei. For each experimental replicate, nuclei were counted from 5 random fields taken with a 10X objective equipped in a NIKON Eclipse 80i fluorescence microscope. For primary mouse myoblast cell count, myoblasts were plated at equal densities following quantification with a hemocytometer. For each time point, cells were stained with DAPI. Each experimental replicate was quantified as the average from 5 random fields. Individuals taking pictures and counting nuclei were blinded to genotype.

### Western blot

Muscle tissues and cultured cells were homogenized using RIPA lysis buffer (50 mM Tris–HCl, pH 7.4, 150 mM NaCl, 1% NP-40, 0.25% Na-deoxycholate, and 1 mM EDTA) supplemented with protease inhibitor cocktail (Millipore Sigma), and lysates were centrifuged at 16,000*g* for 10 minutes. Protein concentrations were determined using a BCA Protein Assay Kit (Thermo Fisher Scientific). Aliquots of lysates were solubilized in Laemmli sample buffer, and equal amounts of proteins were separated on 12% SDS-PAGE gels. Proteins were transferred onto nitrocellulose membranes and then blocked with 5% nonfat dry milk in PBS with 0.1% Tween-20 for 1 hour. The membrane was then incubated with primary antibodies, specific to the protein of interest, in 5% nonfat dry milk in PBS with 0.1% Tween overnight at 4°C. After incubation with the appropriate secondary antibody conjugated with HRP, ECL (GH Healthcare) was used for protein detection. Immunoblots were obtained using the G:Box Chemi XT4, Genesys, version

1.1.2.0 (Syngene). Densitometry was measured with ImageJ software (NIH).

### Antibodies

The antibodies used were as follows: anti–rabbit GAPDH (Cell Signaling Technology; 2118), anti–mouse desmin (Dako; M0760), anti–rabbit DNAJB6 (Abcam; ab198995), anti–mouse HSP70 (Enzo Life Sciences; ADI-SPA-812) anti-rabbit Synemin (Bioss, bs-8555R), anti-rabbit alpha Actinin (Abcam ab68167), anti-rabbit Cathepsin-D (Abcam, ab6313), anti-mouse myosin (Sigma, M1570), anti-rabbit DNAJB4 (Proteintech, 13064-1-AP), and anti-rabbit FHL2 (Abcam ab202584). The following secondary antibodies were used: anti–mouse HRP (Cell Signaling Technology; 7076S), anti–rabbit HRP (Cell Signaling Technology; 7074S), and anti–rabbit Alexa Fluor 555 (Invitrogen, Thermo Fisher Scientific).

### Real-Time qPCR

Total RNA was isolated from primary myotubes with the SV Total RNA Isolation Kit (Promega; Z3100) according to the manufacturer’s instructions. The concentration and quality of the total RNA isolated were determined using a NanoDrop spectrophotometer (Thermo Fisher Scientific). cDNA was synthesized using the Transcriptor First-Strand cDNA Synthesis Kit with anchored oligo (dT)_18_ primers (Roche; 04379012001). Gene expression levels were analyzed by real-time PCR on an Applied Biosystems model 7500 Software (version 2.0.5) using FastStart Universal SYBR Green Master ROX qPCR Mastermix (Roche; 04913850001). qPCR was performed with primers for GAPDH (F: ATGGTGAAGGTCGGTGTGA, R: AATCTCCACTTTGCCACTGC), DNAJB6a (F: TGGGTCTAAAAGCAACTGGG, R: AGTCCTCTTTCTGCTTCTGC), and DNAJB6b (F: ACGACAAAGAGGATTGTGGAG, R: GCAGGTGCTCCTTACCATTTA). Values were normalized to GAPDH and are presented in log scale as -ddCT. Each data point (experimental replicate) was generated from three technical replicates.

### TMT Labeling Proteomics

Primary myotubes were grown and treated with morpholinos as described above. Cell pellets were prepared for mass spectrometry acquisition by following the EasyPep Mini MS Sample Prep Kit (Thermo Fisher Scientific Inc.). Peptide concentrations were determined with the Quantitative Fluorometric Peptide Assay (Thermo Fisher Scientific Inc). A total of 10μg of peptide was prepared to label with TMTpro 16 plex reagents (Thermo Fisher Scientific Inc.) according to the manufacturer’s instructions. Labeled samples were combined and dried with vacuum centrifugation. Samples were then separated into eight fractions using the High pH Reversed-Phase Peptide Fractionation Kit (Thermo Fisher Scientific Inc.). The fractions were dissolved in 0.1% formic acid, and peptide concentrations were determined with Quantitative Colorimetric Peptide Assay (Thermo Fisher Scientific Inc.). TMT labeled sample LC-MS/MS acquisitions were performed using an EASY-nLC 1000 connected to an Orbitrap Eclipse Tribrid mass spectrometer (Thermo Fisher Scientific Inc.). An amount of 0.5ug of each fraction was loaded on an Aurora UHPLC Column (Ionopticks, Fitzroy, Australia) and separated with a 136-min method, as described previously.^30^ MS1 scans were acquired in the Orbitrap at 120 k resolution with a scan range of 350–1800 m/z. The AGC target was 1×10^6^, and the maximum injection time was 50 ms. MS2 scans were acquired with higher-energy collisional dissociation (HCD) activation type in the Orbitrap at 50 k resolution with the defined first mass 110 m/z. The isolation window was 0.5 m/z, collision energy was 38%, maximum injection time was dynamic, and AGC target was standard. System control and data collection were performed with Xcalibur software (Thermo Fisher Scientific Inc.).

Proteomic analyses were performed with Proteome Discoverer 2.4 (Thermo Fisher Scientific Inc.) using the Uniprot human database and the SequestHT with Percolator validation. TMTpro (Any N-Terminus) was set as a static N-Terminal Modification; TMTpro (K) and carbamidomethyl (C) were set as static modifications; oxidation (M) was set as a dynamic modification; acetyl (protein N-term), Met-loss (Protein N-term M), and Met-loss + acetyl (Protein N-term M) were set as dynamic N-Terminal modifications. Normalization was performed relative to the total peptide amount. Further analyses were performed using the normalized abundance as below: limma analyses were performed using R studio following the user guide;^31^ and volcano plots and heatmaps were generated with Prism 9.

Proteins were classified broadly into several catalogs according to the Gene Ontology (GO) annotation (geneontology.org). Overrepresentation analyses of GO terms, including biological process, molecular function, and cellular component, were performed using the ConsensusPathDB-human database system (cpdb.molgen.mpg.de/CPDB), which is a molecular functional interaction database. All proteins detected in the mass-spectrometry experiments were used as background for comparison. A p-value cutoff of 0.01 was selected.

### Statistics

Comparisons between 2 groups were made using a 2-tailed, unpaired Student’s *t* test. Comparisons between several groups were made using either a 1-way ANOVA with Tukey’s post hoc test or two-way ANOVA with Bonferroni’s post hoc test to adjust for multiple comparisons. All analyses were performed with GraphPad Prism 9 (GraphPad Software). Results were considered statistically significant if *P* was less than 0.05.

## Acknowlegment

This work was supported by grants from: The National Institute of Arthritis and Musculoskeletal and Skin Diseases (NIAMS): ARF (K08AR075894, R03AR081395), CCW (R01AR068797, K24AR073317); The American Society of Gene and Cell therapy (ASGCT): ARF (Career Development Award); The Children’s Discovery Institute of Saint Louis Children’s Hospital: ARF (Faculty Scholar Award-MIFR20221004); and the LGMD-1D DNAJB6 Foundation and International Registry (ARF).

## Author Contributions

ARF: Conceptualization, formal analysis, funding acquisition, investigation, methodology, resources, supervision, validation, visualization, writing original draft, writing – review and editing.

MMP, JAD, RB, SKP, SL, FW, TC: Investigation, methodology.

HLT, TMM: Conceptualization, writing – review and editing.

CCW: Conceptualization, funding acquisition, resources, supervision, validation, writing – review and editing.

## eTOC synopsis

Dominant point mutations in *DNAJB6* cause LGMDD1. Knockout is embryonic lethal. Selective knockdown treatments may improve therapeutic index. Findlay and colleagues used morpholinos to knockdown DNAJB6b, an isoform central to pathogenesis, and correct disease phenotypes in models of LGMDD1. Selective DNAJB6b knockdown may be a viable therapeutic target for LGMDD1.

